# CHDbase: A Comprehensive Knowledgebase for Congenital Heart Disease-related Genes and Clinical Manifestations

**DOI:** 10.1101/2022.01.06.475217

**Authors:** Wei-Zhen Zhou, Wenke Li, Huayan Shen, Ruby W. Wang, Wen Chen, Yujing Zhang, Qingyi Zeng, Hao Wang, Meng Yuan, Ziyi Zeng, Jinhui Cui, Chuan-Yun Li, Fred Y. Ye, Zhou Zhou

**Author notes:** **Corresponding authors.** (Zhou Z), (Ye FY). Equal contribution (Wei-Zhen Zhou, Wenke Li, Huayan Shen).

## Abstract

Congenital heart disease (CHD) is the most common cause of major birth defects, with a prevalence of 1%. Although an increasing number of studies reporting the etiology of CHD, the findings scattered throughout the literature are difficult to retrieve and utilize in research and clinical practice. We therefore developed CHDbase, an evidence-based knowledgebase with CHD-related genes and clinical manifestations manually curated from 1114 publications, linking 1124 susceptibility genes and 3591 variations to more than 300 CHD types and related syndromes. Metadata such as the information of each publication and the selected population and samples, the strategy of studies, and the major findings of study were integrated with each item of research record. We also integrated functional annotations through parsing ~50 databases/tools to facilitate the interpretation of these genes and variations in disease pathogenicity. We further prioritized the significance of these CHD-related genes with a gene interaction network approach, and extracted a core CHD sub-network with 163 genes. The clear genetic landscape of CHD enables the phenotype classification based on the shared genetic origin. Overall, CHDbase provides a comprehensive and freely available resource to study CHD susceptibility, supporting a wide range of users in the scientific and medical communities. CHDbase is accessible at http://chddb.fwgenetics.org/.

## Introduction

Congenital heart disease (CHD) refers to abnormalities in cardiocirculatory structure or function that arise during embryonic development. With a prevalence as high as ~1% of live births [1], CHD is the most common cause of major birth defects [2] and imposes enormous health and economic burdens. Specifically, 3%–5% of CHD patients have a genetic syndrome manifesting not only as CHD but also extracardiac defects, which is termed syndromic CHD; others present abnormalities restricted to the heart, which is termed nonsyndromic CHD. Family studies have shown the strong genetic susceptibilities underlying the pathogenesis of CHD [3, 4], it is thus essential to clarify the CHD genetic etiology in the diagnosis and treatment of CHD. Recently, numerous genetics studies have addressed the genetic susceptibilities underlying CHD. Briefly, it has been estimated that the chromosomal aberrations and aneuploidies account for 8%–10% of CHD, the copy number variations (CNVs) for 3%–10% of CHD [5], *de novo* and rare inherited variants for 8% and 2% of CHD, respectively [6].

However, as the findings of these genetics studies are largely scattered in the literature, it is difficult to incorporate them in the in-depth mechanism studies and clinical practice. The holistic picture of CHD susceptibility, such as the number and features of CHD-related genes, the strength of each item of evidence, and the relationship between genetic elements and cardiac malformations, is still to be addressed. Notably, the adequate classification of CHD types could substantially facilitate the investigation of its pathogenesis, the accurate diagnosis, and the effective treatments, while the current practice largely depending on its clinical manifestations with high diversity [7–9]. Efficient molecular classification could certainly supplement the current practices, while it is still to be addressed due to the lack of well-organized genotype–phenotype correlations. Notably, one pilot study recently reported a CHD-RF-KB database [10] to curate genetic risk factors associated with nonsyndromic CHD from approximately 300 studies, while it is far from complete to provide a clear genetic landscape of CHD for basic and translational studies.

Here, we present a comprehensive knowledgebase for CHD with extensive research evidence and functional annotations. We also prioritized these CHD-related genes with a gene interaction network approach. We then classified the CHD types based on their shared genetic origin, facilitating a molecular diagnosis and treatment of CHD which supplementing the traditional practices with clinical manifestations.

## Database implementation

### Data collection and processing

To integrate current knowledge of CHD, we searched the PubMed database to retrieve CHD-related publications, using keywords including “congenital heart disease”, “heart defects”, “gene”, “proteomics”, “expression”, “copy number variation”, “linkage”, “association”, and “sequencing” (see supplementary methods in File S1). Among more than 2700 search results, we confirmed the relevance of each publication by examining its title and abstract. We excluded publications about congenital cardiomyopathies or congenital heart rhythm disorders due to their distinct clinical presentation and the underlying mechanisms. After this process, a total of 1114 research articles published between February 1992 and January 2020 were collected for further curation.

For each article, we reviewed the main text and additional materials to extract evidence of association between genes/variations and CHD. If multiple experimental approaches or different datasets were used in one study, the evidence of this study was divided into multiple independent records. In total, 1353 items of evidence were extracted, which could be classified into six types as follows: 1) 159 items of “Genetic association” evidence, 2) 452 items of “SNV/Indel” evidence reporting single-nucleotide variants/insertion-deletion variants (SNVs/Indels) in CHD patients, 3) 63 items of “Expression” evidence linking differentially-expressed genes to CHD, 4) 240 items of “CNV” evidence reporting CNVs detected in patients, 5) 15 items of “Linkage” evidence reporting CHD-related genomic regions by linkage studies, and 6) 424 items of “Other” evidence, such as the evidence from transgenic animal models and cell lines (**Figure 1**). For each item of evidence, comprehensive metadata, such as the information of each publication and the selected population and samples, the strategy of studies, and the major findings of association, were extracted manually and double-checked by two researchers (Figure 1). A detailed description of the collected information is listed in Table S1.

**Figure 1.**
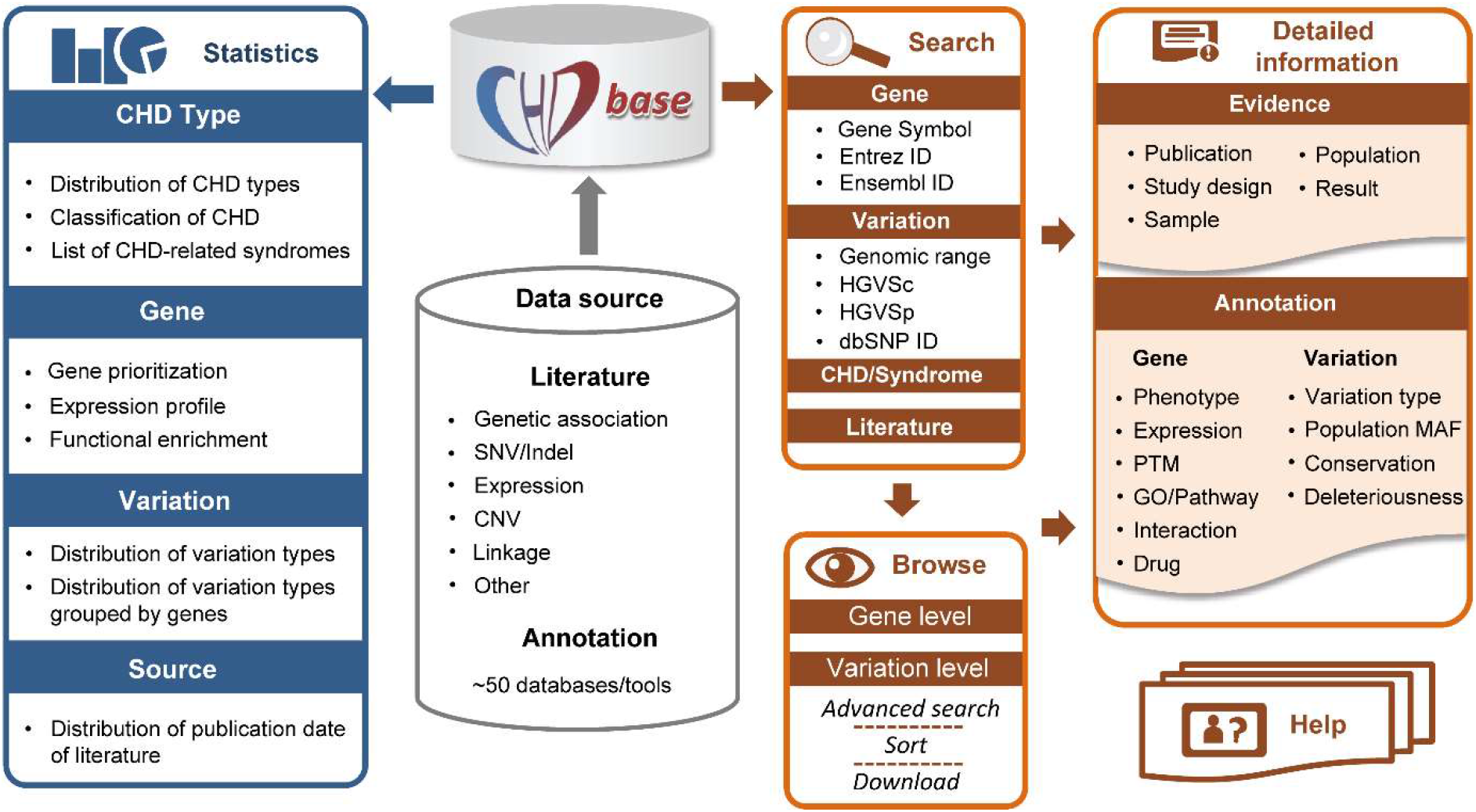
Overview of the framework of CHDbase. PTM, posttranslational modification; GO, Gene Ontology; MAF, minor allele frequency.

The disease name was standardized according to the 11th revision of the International Classification of Diseases (ICD-11) congenital cardiology terms [11]. Transcript-based SNVs/Indels were annotated by Ensembl Variant Effect Predictor (VEP) [12] and confirmed by Mutalyzer [13] following the nomenclature recommendations of the Human Genome Variation Society (HGVS). Genomic coordinates are presented according to the human reference genome (GRCh37/hg19). Overall, CHDbase integrated 3813 gene–CHD associations and 3957 variation–CHD associations, which contains the association information for 1124 susceptibility genes, 1006 structural variations, and 2585 SNVs/Indels.

### Prioritization of CHD-related genes

For gene prioritization, traditional evidence-based prioritization approach typically assigns a gene to a weighted sum based on the number and type of evidence. As the weight for each type of evidence is debatable, and the items of evidence from literature may simply reflect hot research topics rather than the significance of each gene, here we introduced a gene interaction network approach to prioritize the significance of CHD-related genes.

In CHDbase, 1124 genes have been linked to CHD susceptibility, including 115 reported only in syndromic CHD (syndromic genes), 597 reported only in nonsyndromic CHD (nonsyndromic genes), and 412 reported in both syndromic and nonsyndromic CHD (Figure S1). We further prioritized these susceptibility genes with a gene interaction network approach and extracted a core CHD gene set (see supplementary methods in File S1). Briefly, on the basis of the 1124 susceptibility genes and protein–protein interaction annotations from the STRING database [14], we first constructed an unweighted gene interaction network with 952 nodes and 30,777 edges. Then, we used three centrality scores, namely, degree, betweenness centrality, and eigenvector centrality, to measure the significance of each gene in the network. Specifically, a gene with a high degree could be considered as a “hub” in the CHD network with more interactions with other genes. A gene with large betweenness centrality could be considered as a “bridge” in the network and the removal of such a gene could lead to a disconnected network. A gene with large eigenvector centrality has more interactions with high-degree genes. A gene with high degree, large betweenness centrality, and large eigenvector centrality could be more important within the network.

To obtain the core gene set, we extracted a k-core with 163 genes from the CHD network using k-core decomposition (see supplementary methods in File S1). As we retained the nodes with a maximum value of *k (k* = 70), these 163 genes in the k-core were considered as the most centrally located nodes in the original network. The core gene set contains six syndromic genes, 83 nonsyndromic genes and 74 genes reported in both syndromic and nonsyndromic CHD, indicating that the syndromic and nonsyndromic CHD share a common gene network (**Figure 2**A). These core genes are supported by a significantly higher number of supporting evidence than that of all CHD-related genes (*P* = 4.55E-10, Mann–Whitney U test). Notably, for 38 high-confidence CHD genes recurrently reported in recent reviews [5, 15–18], 18 are present in this core gene set (Figure 2B). To facilitate users to rank genes based on their own experiences and preferences, we provide the CHD susceptibility gene list with the meta-data, such as the information indicating whether it belongs to the core gene set, its centrality scores, and the number of evidence items supporting its association with CHD, as shown in Table S2.

**Figure 2.**
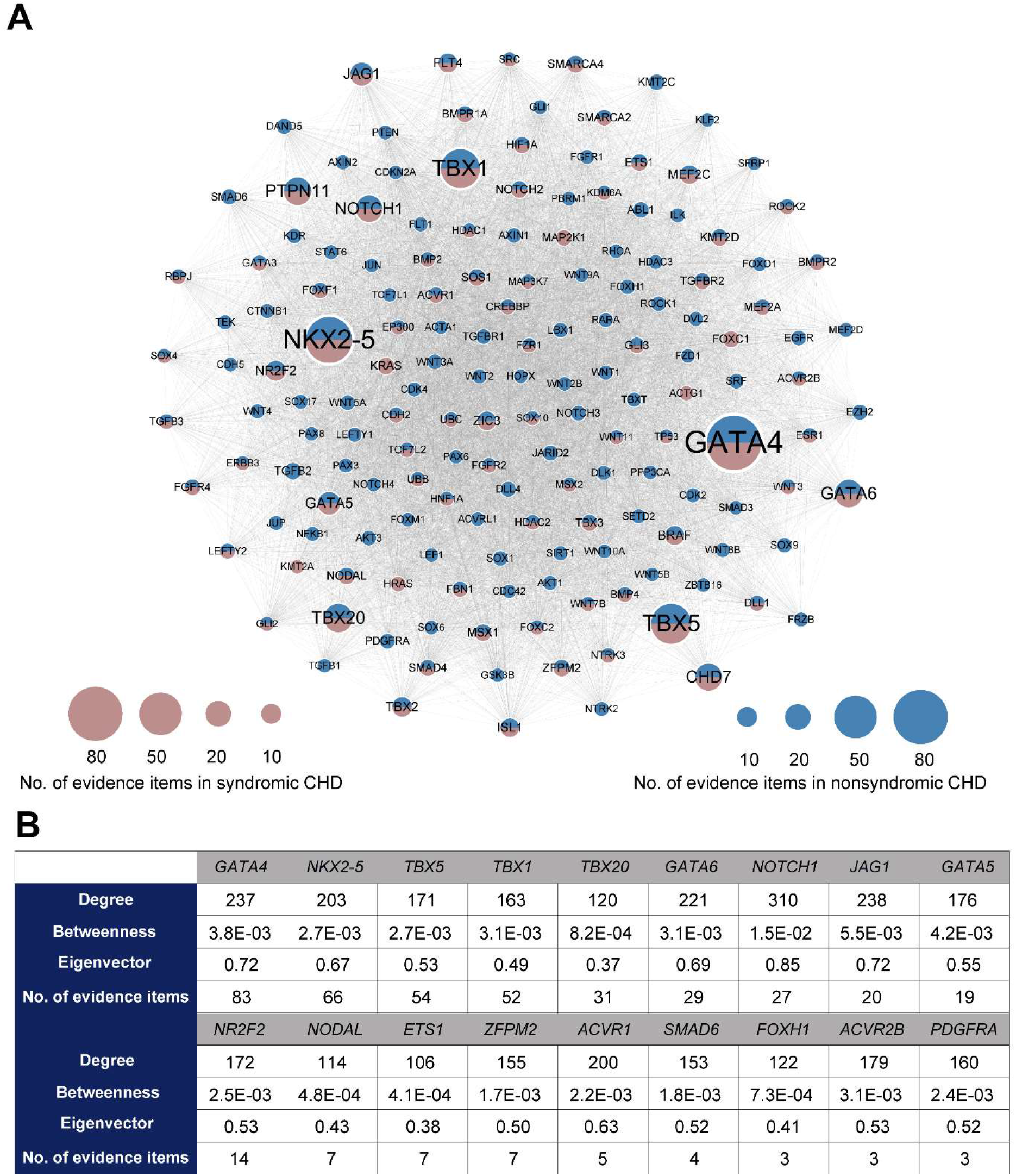
Network-based prioritization of CHD-related genes. **A.** The k-core of CHD-related genes (n = 163). The node size is proportional to the number of supporting evidence items for each gene. **B.** For 18 core genes overlapping with the high-confidence CHD genes recurrently reported in recent reviews, the degree, betweenness centrality, eigenvector centrality, and the number of supporting evidence items for each gene are listed.

### Functional annotations of CHD-related genes

To better understand the function of CHD-related genes, we further annotated them using public databases and data as follows: i) HGNC (https://www.genenames.org/), NCBI Entrez gene [19], Ensembl (http://www.ensembl.org/), Genecards [20], OMIM [21], UniProt (http://www.uniprot.org/), Mouse Genome Informatics (MGI) [22], and Zebrafish Model Organism Database (ZFIN) [23] for basic information of the gene and its orthologs; ii) possible associated phenotypes and diseases from Human Phenotype Ontology (HPO) [24], GWAS Catalog [25], and DisGeNET [26]; iii) spatiotemporal expression profiles integrated from the genotype-tissue expression (GTEx) project [27], RNA-Seq data for seven human organs across multiple developmental time points [28], and spatial subcellular expression data of the developing human heart at three developmental phases [29]; iv) posttranslational modifications retrieved from UniProt and DbPTM [30]; v) biological pathways and protein–protein interactions annotated by Gene Ontology (GO, http://www.geneontology.org/), Reactome [31], BioCyc [32], KEGG [33], PANTHER [34], STRING [14], OmniPath [35], BioGRID [36], and the human reference interactome (HuRI) database [37]; and vi) drug-gene interactions retrieved from DGIdb [38], DrugCentral [39], and PharmGKB [40] (Figure 1). Additionally, to better understand the effects of variations, we classified the variations into five categories according to the conclusions of the original publications: “disease causing”, “likely disease causing”, “uncertain”, “likely benign” and “benign”. For SNVs/Indels, we further integrated genomic information annotated by VEP [12], including variant consequences, minor allele frequency in different populations of gnomAD [41], conservation scores calculated with phastCons, phyloP, and GERP++, and variant deleteriousness predicted by SIFT, PolyPhen, CADD, DANN, LRT, M-CAP, MetaLR, PrimateAI, fathmm-MKL, and fathmm-XF (Figure 1).

### Database interface

We constructed a MySQL database to store and manage the data. A user-friendly web interface was further implemented in Java, HTML, CSS, and JavaScript, as powered by SpringBoot, Thymeleaf, and Bootstrap framework.

### User interface and functions

CHDbase provides two search modes. The basic search mode at the Home Page and the top navigation bar allows users to precisely or fuzzily search for genes, variations, CHD/syndrome and literature of interest (**Figure 3**A, B). The basic search engine can recognize query terms such as the gene symbols, Entrez IDs, Ensembl IDs, the range of genomic coordinates, HGVS expressions at the cDNA level (HGVSc), HGVS expressions at the protein level (HGVSp), dbSNP IDs, the full name or abbreviation of diseases, and PubMed IDs. When the user clicks on the button of “Exact Search”, a result page fully matching the search term displays the detailed evidence of gene– or variation–CHD association. If the user clicks on the “Fuzzy Search” button, the search engine will attempt to find partial matches for search term, and return the results on the Browse Page. The Browse Page not only enables users to flexibly browse associations at the gene level or variation level, but enables the users to filter data with an advanced search mode (Figure 3C). Specifically, the user can enter multiple search terms in corresponding search boxes under column names to search for information of interest. Search results can be sorted by multiple columns in a customized order, and are downloadable in JSON or CSV format. The user could further click on the gene or variation links on the Browse Page to view the detailed information of research evidence and annotations as described below.

**Figure 3.**
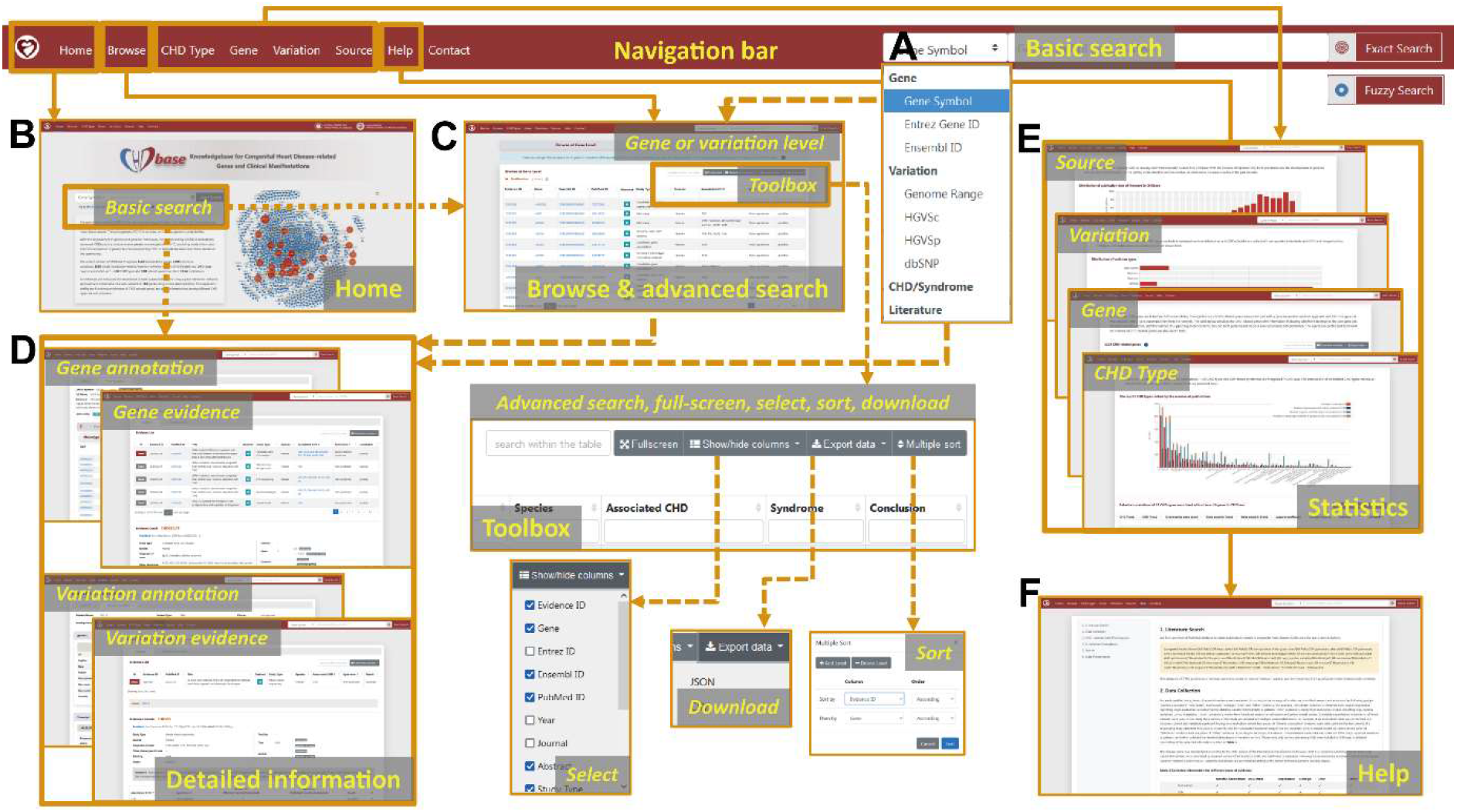
The web interface of CHDbase. The major components for the web interface of CHDbase are shown, including the navigation bar (**A**), the Home Page (**B**), the Browse Page (**C**), and the Evidence and Annotation Page for each CHD-related gene or variation (**D**). The Statistics Page for the diseases, genes, variations, and studies (**E**), as well as the Help Page (**F**) are also available on the web interface.

Detailed information about gene–CHD or variation–CHD associations is provided in a similar manner on the Gene or Variation Evidence Page (Figure 3D). Taking Gene Evidence Page as an example (Figure S2), in the upper panel, the list of research evidence for a specific gene relevant to CHD is summarized, including evidence ID, PubMed ID, title, abstract, study type, species, associated CHD types, associated syndromes, and conclusions. Furthermore, the user can click on the “Detail” label on the left of the “Evidence ID” column to view details in the lower panel. For detailed information on each item of evidence, we first provide population information, study design, and a summary of results, followed by the details of the results. According to different categories of evidence, the detailed information is summarized in varied formats (Figure S3). For example, for the evidence of “SNV/Indel” or “CNV”, variation, variation type, associated CHD types, associated syndromes, No. of variation in affected individuals, No. of variation in unaffected individuals, and variation pathogenicity are provided. If the sample information is available, the user can click on the plus sign at the end of the row to view the information of individuals carrying this variation, including sample ID, family name, ethnicity, sex, age, diagnosis, other phenotypes, transmission (*de novo,* paternal, maternal, and familial), and zygosity (homozygous, heterozygous, and mosaic). For the evidence of “Genetic association”, “Expression” and “Linkage”, besides the associated CHD types and syndromes, statistical methods and results are also integrated. For further functional information about the gene or variation, the user can click on the label of “Gene Annotation” or “Variation Annotation” in the top left to view annotations from external databases and the data mentioned above (Figure S2, Figure S4).

Finally, to display a comprehensive genetic landscape of CHD, the database also shows the statistics at disease, gene, variation, and study levels, in charts or forms with embedded hyperlinks (Figures 1, 3E). A Help Page with detailed introductions about the database is also provided (Figures 1, 3F).

### Features of CHD-related genes

To clarify the molecular mechanisms of CHD and facilitate the prioritization of novel susceptibility genes, we systematically characterized the features of CHD-related genes. As syndromic CHD is usually accompany with abnormal brain development, we explored the expression profiles of three CHD gene categories in the brain and heart, across different time points of human development [28]. Compared with a random gene set unrelated to CHD as a background, all of the three CHD categories showed higher expression levels in the brain and heart, especially in fetal organs, indicating their roles in embryonic development. Notably, the syndromic genes showed higher expression levels than nonsyndromic genes in the brain (*P* = 0.13, one-sided Mann–Whitney U test) (**Figure 4**A). However, in the heart, nonsyndromic genes expressed at a significantly higher level than syndromic genes (P < 2.20E-16, one-sided Mann–Whitney U test), suggesting a more specific role of nonsyndromic genes in heart development (Figure 4B). Furthermore, CHD-related genes tend to broadly-expressed across tissues and developmental stages compared with the background, according to tissue- and time-specificity indexes as defined by Cardoso-Moreira *et al.,* indicating CHD-related genes are more intolerant to functional variations [28] (Figure 4C, D).

**Figure 4.**
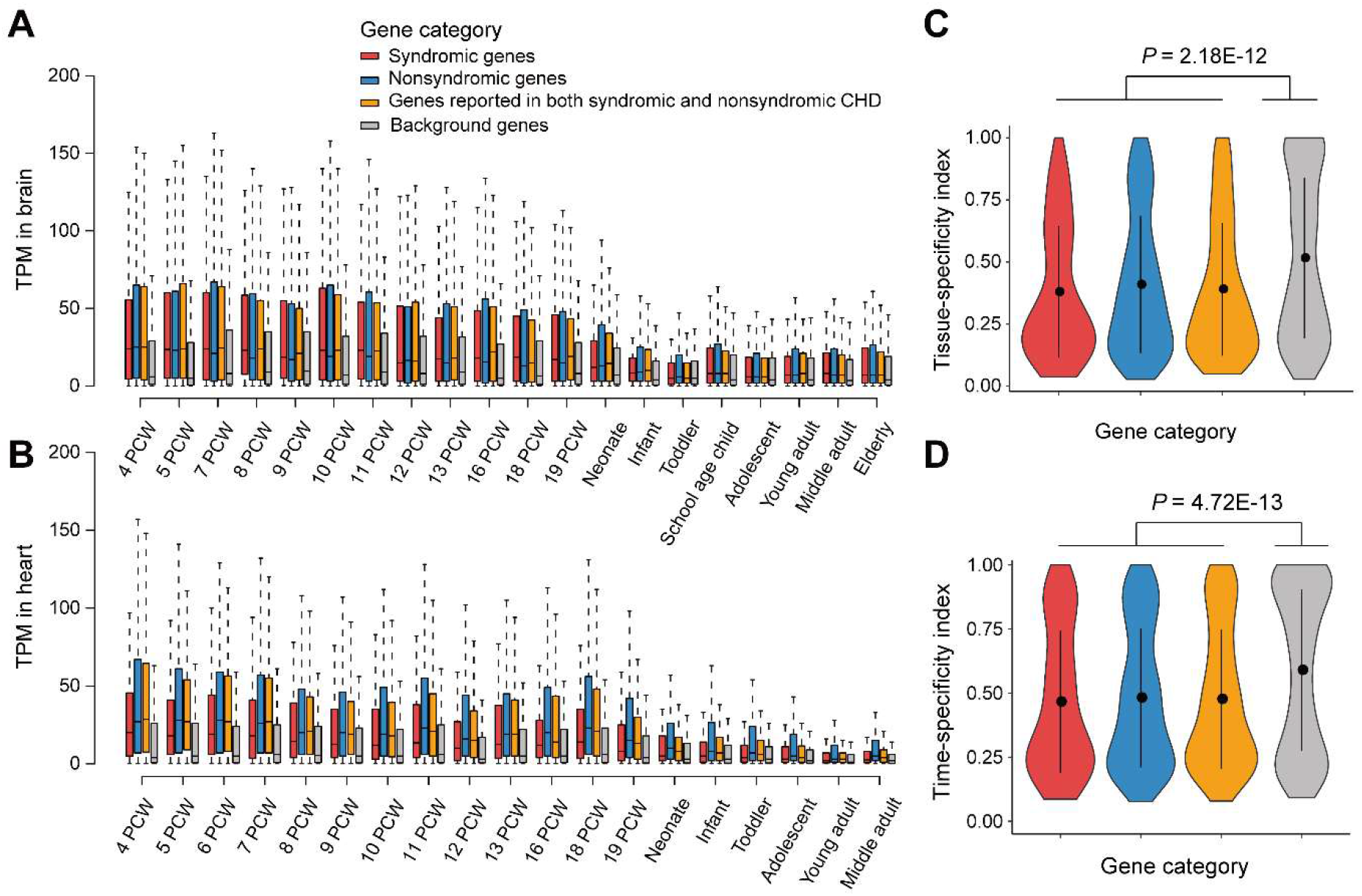
The expression profiles of CHD-related genes. Boxplot showing the expression levels for three categories of CHD-related genes in the brain (**A**) and heart (**B**) across different developmental time points. Violin plot showing the difference between CHD-related genes and background genes in tissue-specificity (**C**) and time-specificity (**D**). The P values were obtained with Mann–Whitney U test. In above figures, CHD-related genes are classified into three categories: syndromic genes (red, n = 115), nonsyndromic genes (blue, n = 597), and genes reported in both syndromic and nonsyndromic CHD (orange, n = 414). A non-CHD gene set (n = 1124) was randomly selected as the background (gray). TPM, transcripts per million; PCW, post conception week.

We then identified the enriched GO terms and pathways for CHD-related genes with ClueGO [42] (**Figure 5**, Table S3). Briefly, the GO terms of heart development, embryonic morphogenesis, pattern specification process, cell differentiation, muscle structure development, regulation of cellular component movement and cell surface receptor signaling pathway are significantly enriched. Consistently, on pathway level, the cell surface receptor signaling pathways involved in heart development, such as the Wnt, Notch, and TGF-beta signaling pathways, are significantly enriched. The enrichment of CHD-related genes on terms of heart development is expected, since the initial keywords for retrieving CHD-related genes were chosen to find these genes. While the enriched terms and pathways identified here provide a comprehensive molecular picture to understand CHD. Moreover, several cancer-related signaling pathways are also identified, providing a perspective to understand the previous epidemiological data that CHD patients typically show higher cancer prevalence [43, 44]. Interestingly, the generic transcription pathway is also enriched, with 112 of the 1124 CHD-related genes cataloged as transcription factors in hTFtarget [45] (*P* < 2.20E-16, Fisher’s exact test). Notably, among the top ten genes with the strongest supporting evidence, six are transcription factors (*GATA4, NKX2-5, TBX5, GATA6, CHD7,* and *NOTCH1*). In addition, our data also recapitulated several pathways previously linked to CHD etiology [18], such as muscle contraction, extracellular matrix organization and cilium assembly.

**Figure 5.**
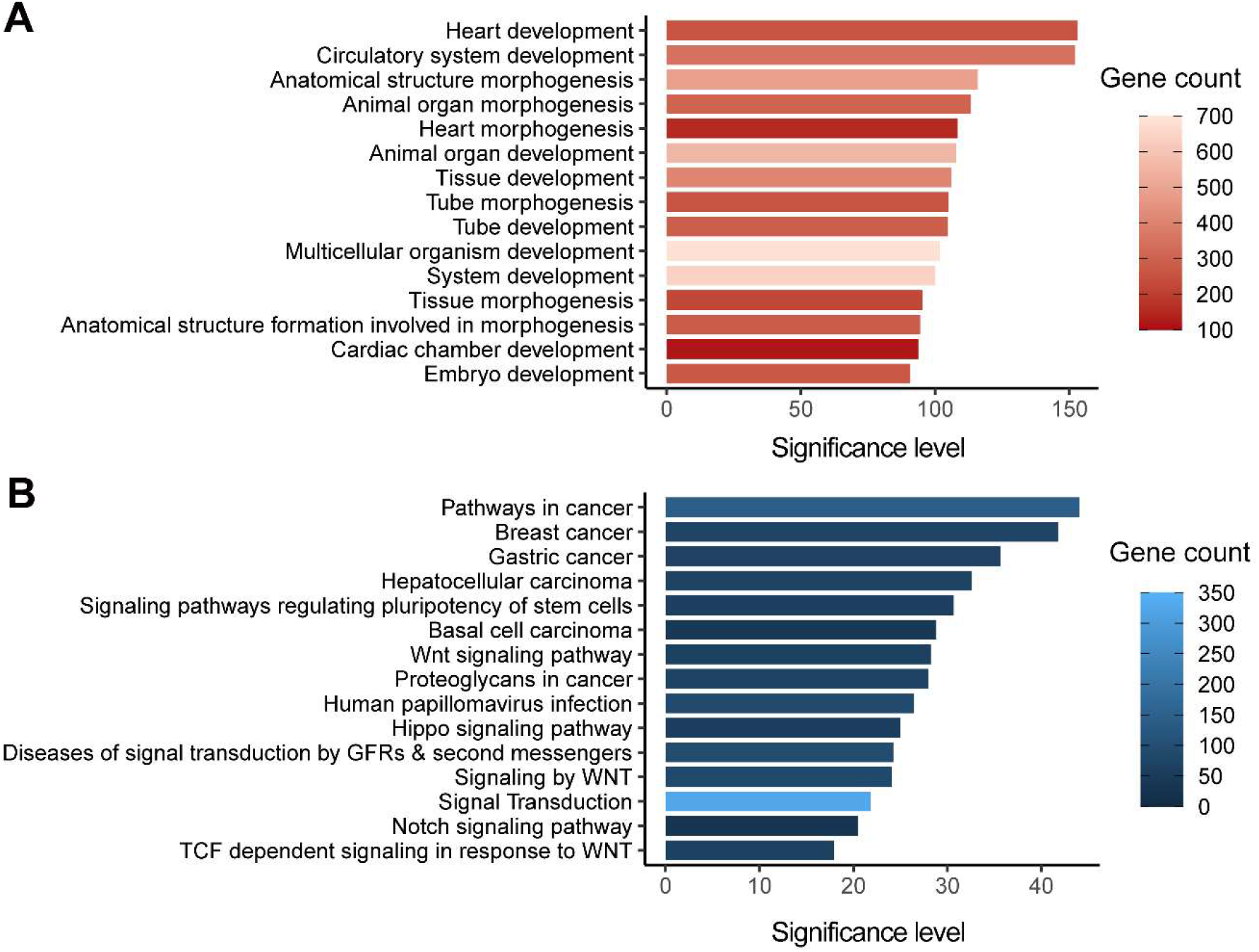
Functional enrichments of CHD-related genes. **A.** The top 15 significantly enriched Gene Ontology terms for CHD-related genes (n = 1124) are shown with significance levels. **B**. The top 15 significantly enriched KEGG and Reactome pathways for CHD-related genes (n = 1124) are shown with significance levels. The adjusted *P* values corrected with Bonferroni step down were -log (base 10) transformed to denote the significance levels. GFR, growth factor receptor.

### Molecular classification of CHD

CHD is a highly heterogeneous disease with a large set of cardiac malformations. Adequately clustering these malformations could classify this complex disease into several groups with shared pathophysiological mechanism, facilitating in-depth mechanism study and clinical management. As CHDbase integrates comprehensive phenotype–genotype correlation data, we then attempted to classify CHD based on the genetic background (see supplementary methods in File S1). We first used Jaccard coefficient [26] to measure pairwise genetic similarity between all pairs of 27 CHD types. In total, 196 pairs of CHD types showed significantly more shared genes than expected (adjusted P values < 0.05) (**Figure 6**A, Table S4). Based on the matrix of pairwise Jaccard distances, hierarchical clustering was further performed to reveal genetic patterns shared by CHD types. Seven distinct groups were identified (Figure 6B), which are largely consistent with known developmental processes and previous CHD classifications, such as Botto’s taxonomy [7] and Ellesøe’s classification based on familial co-occurrence [9]. To control for the effects of false positives in identifying disease-causing variants, we repeated the classification after excluding 21 variants that were reclassified as “benign” or “likely benign” by InterVar [46] according to the American College of Medical Genetics and Genomics/Association for Molecular Pathology (ACMG/AMP) guidelines [47]. The new results are consistent, indicating the reliability of this classification.

**Figure 6.**
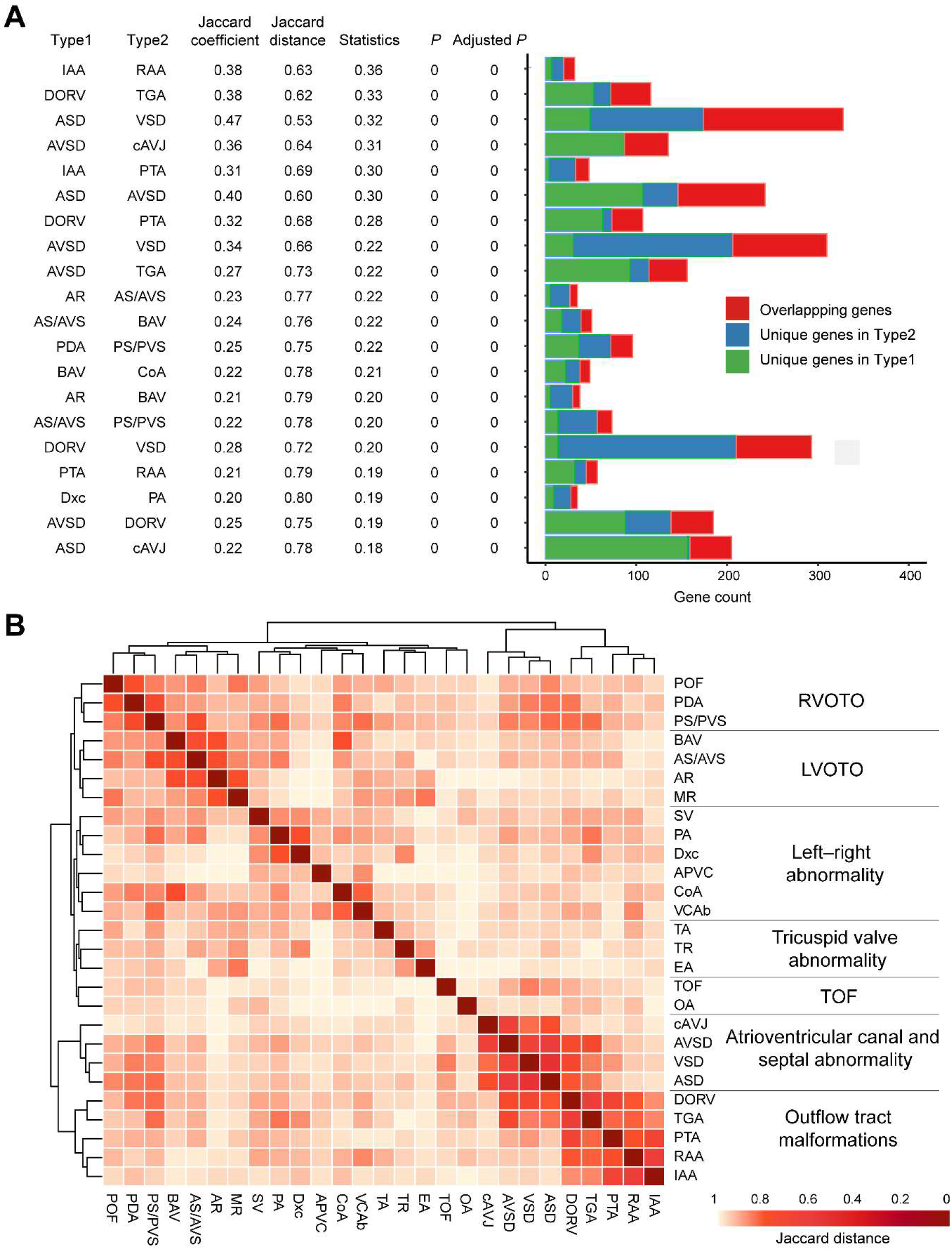
CHD classification based on genotype–phenotype correlations. **A.** Jaccard coefficient, Jaccard distance, statistical significance for Jaccard coefficient, and gene distribution are shown for 20 pairs of CHD types with the strongest correlations. Statistics: the centered Jaccard coefficient, which equals to the Jaccard coefficient minus the unbiased estimation of expectation. P and Adjusted P: calculated with the Jaccard test and corrected for multiple testing using the Benjamini–Hochberg false discovery rate. **B.** Classification of 27 CHD types in CHDbase. Only CHD types associated with at least ten genes were included in this analysis. Jaccard distances for all pairs of CHD types are shown in a heatmap, which were further used to classify these CHD types into seven major groups through hierarchical cluster analysis. For each group, the anatomical term is defined and shown. POF, patent oval foramen; PDA, patent arterial duct; PS/PVS, pulmonary stenosis/pulmonary valve stenosis; BAV, bicuspid aortic valve; AS/AVS, aortic stenosis/aortic valvar stenosis; AR, aortic regurgitation; MR, mitral regurgitation; SV, single ventricle; PA, pulmonary atresia; Dxc, dextrocardia; APVC, anomalous pulmonary venous connection; CoA, coarctation of the aorta; VCAb, vena cava abnormality; TA, tricuspid atresia; TR, tricuspid regurgitation; EA, Ebstein malformation of the tricuspid valve; TOF, tetralogy of Fallot; OA, overriding aorta; cAVJ, common atrioventricular junction; AVSD, atrioventricular septal defect; VSD, ventricular septal defect; ASD, atrial septal defect; DORV, double-outlet right ventricle; TGA, transposition of the great arteries; PTA, persistent truncus arteriosus; RAA, right aortic arch; IAA, interrupted aortic arch; RVOTO, right ventricular outflow tract obstruction; LVOTO, left ventricular outflow tract obstruction.

As depicted in Figure 6B, cardiac malformations of abnormal rotation and septation of the outflow tract, such as transposition of the great arteries (TGA), double-outlet right ventricle (DORV), persistent truncus arteriosus (PTA), interrupted aortic arch (IAA), and right aortic arch (RAA), were grouped into “Outflow tract malformations”, which represents the spectrum of conotruncal defects. This group highly correlates with “Atrioventricular canal and septal abnormality” because it is usually accompanied by septal abnormalities. In addition to “Outflow tract malformations”, ventricular outflow tract obstruction (VOTO) clustered strongly and was divided into left (LVOTO) and right (RVOTO) groups. VOTO refers to any anatomic or functional obstruction of flow out of the ventricle into the aorta or the pulmonary artery, usually caused by valvular stenosis or atresia. Our LVOTO and RVOTO groups largely overlap with Botto’s and Ellesøe’s classifications. Exceptions are pulmonary atresia (PA) and coarctation of the aorta (CoA). In our classification, PA and CoA share the most genes with dextrocardia (Jaccard coefficient=0.20, adjusted P=0) and vena cava abnormalities (Jaccard coefficient=0.18, adjusted P=0), respectively (Figure 6A); both classified as “left-right abnormality”. As previously reported, PA manifests in 23.2% of patients with heterotaxy, an abnormality of formation of the left-right axis of the body [48], and CoA has been associated with an isolated persistent left superior vena cava, a common vena cava abnormality in 21.3% of patients [49]. These findings support the correlation between the development of these two cardiac malformations and abnormal left–right arrangement. The other four CHD types in our “left–right anomaly” group have been recognized to be related to the left–right arrangement [7, 9]. Taken together, these major groups we defined are highly related to anatomy, embryology, and especially to the genetic origin. This molecular classification would thus serve as a supplement to the traditional CHD classification and clinical practice.

### Comparison with existing databases

To evaluate the utility and uniqueness of CHDbase, we compared it with CHD-RF-KB [10], a recently published database to compile risk factors associated with nonsyndromic CHD but not syndromic CHD, and two widely-used variant databases, ClinVar [50] and the Human Gene Mutation Database (HGMD) [51], that focus on a wide range of inherited diseases. As shown in **Table 1,** CHDbase compiles a larger set of CHD-related genes and variations, from a wider scope and a larger number of publications, compared with the other three databases. In comparison to the three databases which only reported annotations at the variation level, CHDbase provides 60 types of metadata, such as the study design, major associations, and the sample information (Table S1). Moreover, CHDbase provides comprehensive functional annotations and analyses on the genes, showing a comprehensive landscape of CHD susceptibility and facilitating a “one-stop” solution to clarify the gene–CHD associations.

**Table 1.** Comparison with existing databases.

|  | **CHDbase** | **CHD-RF-KB<sup>a</sup>** | **ClinVar<sup>b</sup>** | **HGMD<sup>b</sup>** |
| --- | --- | --- | --- | --- |
| **Data source** | Manually curated | Manually curated | User submitted | Manually curated |
| **Target disease** | Syndromic CHD<br>Nonsyndromic CHD | Nonsyndromic CHD | A wide range of diseases | A wide range of diseases |
| **No. of publications** | 1114 | 305 | 163 | 445 |
| **Evidence type** | Genetic association, SNV/Indel, Expression, CNV, Linkage, and Other | Genetic association, SNV/Indel, CNV, and Methylation | Genetic association and SNV/Indel | Genetic association and SNV/Indel |
| **No. of genes** | 1145<sup>c</sup> | 303 | 53 | 874 |
| **No. of variations** | 2585 SNVs/Indels, 1006 CNVs | 1194 SNVs/Indels, 567 CNVs | 774 SNVs/Indels | 2152 SNVs/Indels |
| **Collected information** | Publication, population, study design, result, and sample information | Publication, result, and sample information | Publication and result information | Publication and result information |
| **Annotation** | Gene annotation and variation annotation | Variation annotation | Variation annotation | None |
| **Data analysis** | Expression profile, functional enrichment, and CHD classification | Functional enrichment | None | None |

<sup>a</sup>The genetic data in CHD-RF-KB is summarized (up to 10 January 2020).
<sup>b</sup>The data from published studies is summarized (up to 10 January 2020).
<sup>c</sup>1145 genes: 1124 CHD susceptibility genes and 21 negative genes.

## Discussion

The genetic heterogeneity of CHD has been increasingly recognized in recent decades. However, several fundamental questions, such as the number of genes associated with CHD, the reliability and reproducibility of each association, are still to be addresses, hindering the applications of current knowledge to etiological studies and clinical practice. In this study, we developed an evidence-based, manually curated knowledgebase for CHD, the CHDbase, linking 1124 susceptibility genes and 3591 variations with ~310 CHD types and related syndromes. We also propose a new network-based approach to prioritize the contribution of each gene to CHD pathogenesis. The users could thus simply select their candidate genes for in-depth mechanism study through our user-friendly data retrieval interfaces. CHDbase thus provides a “one-stop” solution from a comprehensive landscape of CHD susceptibility to promising candidate genes, or even drug targets.

Furthermore, we demonstrated the utility of CHDbase through pattern analyses of genetic and phenotypic diversity in CHD. The expression and functional features of CHD-related genes were systemically analyzed to describe the holistic picture of CHD genetic factors, which should facilitate the study of CHD pathogenesis and the identification of novel susceptibility genes. Based on the most comprehensive genotype–phenotype correlations integrated in CHDbase, we further identified CHD groups with similar genetic origin. Although the classification is based on the genetic background, it recapitulates the traditional classification in that similar malformations could be detected in the same cluster, providing new insight into the genetic backgrounds underlying the pathogenesis and recurrence patterns of CHD. Notably, for this approach of classification, the integrity and reliability of genotype–phenotype correlations are crucial. Currently, although hundreds of genetic loci involved in CHD have been reported and integrated, the list of genes associated with CHD is still not complete and conclusive. Thus, the classification in this study is limited to the 27 most frequent CHD types. The molecular classification of CHD could thus be improved along with more accurate gene–CHD associations identified in future.

Taken together, CHDbase can not only improve research on the etiology and pathogenesis of CHD but also aid in clinical practice for CHD, including diagnosis, genetic counseling and treatment. The database will routinely be updated to keep pace with the research progress of CHD and built into a “one-stop” knowledgebase for exploring CHD susceptibility factors.

## Data availability

CHDbase is publicly available at http://chddb.fwgenetics.org/

## CRediT author statement

**Wei-Zhen Zhou**: Conceptualization, Data curation, Formal analysis, Methodology, Visualization, Writing - original draft, Writing - review & editing, Project administration, Funding acquisition. **Wenke Li**: Software, Methodology, Visualization. **Huayan Shen**: Data curation, Supervision, Writing - original draft. **Ruby W. Wang**: Formal analysis, Methodology, Writing - original draft. **Wen Chen**: Conceptualization, Data curation, Supervision. **Yujing Zhang**: Data curation, Visualization. **Qingyi Zeng**: Data curation. **Hao Wang**: Data curation. **Meng Yuan**: Data curation. **Ziyi Zeng**: Data curation. **Jinhui Cui**: Data curation. **Chuan-Yun Li**: Resources, Writing - review & editing. **Fred Y. Ye**: Supervision, Methodology, Writing - original draft. **Zhou Zhou**: Conceptualization, Project administration, Funding acquisition.

## Competing interest

The authors declare no competing interests.

## Acknowledgments

This work was supported by the Young Scientists Fund of the National Natural Science Foundation of China [31801103] and the Chinese Academy of Medical Sciences (CAMS) Initiative for Innovative Medicine [2016-I2M-1-016]. We acknowledge Dr. Adam Yongxin Ye at Harvard Medical School, Boston Children’s Hospital for his support and contribution to the project, and Dr. Ge Gao at Peking University for assistance with database comparison.

## Supplementary material

**File S1 Supplementary methods**

**Figure S1 Chromosome distribution for CHD-related genes**

The color of each gene represents the type of evidence supporting the association of the gene with CHD. The shape of each gene represents the category of the gene.

**Figure S2 Typical Gene Evidence Page and Annotation Page in CHDbase**

The Gene Evidence Page displays the evidence details at the gene level. Gene annotations can be obtained by clicking on the label “Gene Annotation” in the top left.

**Figure S3 Examples of evidence details at the gene level**

As shown in **Figure S2**, the evidence list is shown in the upper panel of the Gene Evidence

Page, and the details of the evidence are shown in the lower panel. In the lower panel, population information, study design, and result summary are provided at the top for all types of evidence, and result details are provided in the lower part in different formats for different types of evidence.

**Figure S4 Typical Variation Annotation Page in CHDbase**

**Table S1 Collected information for different types of evidence**

**Table S2 1124 CHD-related genes**

**Table S3 Functional enrichment results of 1124 CHD-related genes**

**Table S4 Pairwise correlations of CHD types**

## Notes

http://chddb.fwgenetics.org/

### Competing Interest Statement

The authors have declared no competing interest.

