## Supplementary methods and figures for "CHDbase: A Comprehensive Knowledgebase for Congenital Heart Disease-related Genes and Clinical Manifestations"

#### Literature search

We searched the PubMed database to obtain publications related to CHD using the query term "(congenital heart disease*[All Fields] OR heart defect*[All Fields] OR transposition of the great arteri*[All Fields] OR pulmonary atresia[All Fields] OR pulmonary artery atresia[All Fields] OR Anomalous pulmonary venous*[All Fields] OR Ebstein anomaly[All Fields] OR Epstein anomaly[All Fields]) AND (gene[Title/Abstract] AND (proteomics[Title/Abstract] OR expression[Title/Abstract] OR CNV[Title/Abstract] OR copy number variation[Title/Abstract] OR microarray*[Title/Abstract] OR microdel*[Title/Abstract] OR microdup*[Title/Abstract] OR rearrange*[Title/Abstract] OR linkage[Title/Abstract] OR associa*[Title/Abstract] OR scan[Title/Abstract] OR sequenc*[Title/Abstract])) AND ("1000/01/01"[Date - Publication]: "2020/01/10"[Date - Publication])". The abstracts of 2762 studies retrieved were then carefully reviewed to remove irrelevant papers, and the remaining 1114 studies were subjected to systemic curation.

#### Network-based CHD gene prioritization

We developed an unweighted network for 1124 CHD-related genes based on protein–protein interactions from STRING database (https://string-db.org/) [27]. Only experimentally-verified protein–protein interactions from STRING were used. Specifically, in the network, the two genes were connected with an edge only when the STRING experimental score >0. Next, we used three centralities to measure the significance of each gene in the network, including the degree, the betweenness centrality, and the eigenvector centrality [1]. These parameters were calculated in R package “igraph”.

We then used *k*-core decomposition to define the core nodes of the network, and extracted the core sub-network from a large network [2]. Specifically, *k*-core decomposition is started by removing all nodes with degree $k = 1$ from a network. This process may cause new nodes with degree $k \leq 1$, which are also removed until the degree *k* of all the remaining nodes > 1. The removed nodes and their links during the process of $k = 1$ form the 1-shell. Next, this kind of pruning process is continued by $k = 2$ to extract 2-shell and repeated until all higher-layer shells are extracted and all nodes in the network are removed. In practice, we used the function “coreness()” to identify the *k* scores of each nodes. Then the nodes with the highest *k* score ($k = 70$) were defined as core nodes of the network, and the sub-network of these core nodes was defined as the core sub-network. The network was then visualized with the Cytoscape tool (Version 3.8.2) [3]. To evaluate the confidence of the core genes in the *k*-core, we compared the difference between the core genes and all of the 1124 CHD-related genes in the number of supporting evidence items using a two-sided Mann–Whitney U test.

**Expression profile analysis**

We collected RNA-Seq data for different developmental time points in human brain and heart from the European Bioinformatics Institute (E-MTAB-6814, https://www.ebi.ac.uk, data submitted by Cardoso-Moreira *et al.* [4]). Tissue- and time-specificity indexes were also obtained from the study of Cardoso-Moreira *et al.* [4]. The expression profiles, and tissue-

and time-specificity indexes of different gene categories were identified and compared with Mann–Whitney U test.

**CHD classification**

Copy number variations and linkage regions associated with disease typically cover numerous genes, making it difficult to identify true causal genes. We thus excluded copy number variation and linkage data when performing CHD classification. We further removed SNVs/Indels that were classified as “benign”, “likely benign” and “uncertain” by the original publications, as the authors did not draw a concrete conclusion about the association.

We then calculated the pairwise Jaccard coefficient and statistical significance for the 27 most frequently reported CHD types associated with at least ten genes in CHDbase, using the Jaccard.test function of the Jaccard R package with the measure concentration algorithm. The Jaccard coefficient is defined as the number of shared genes divided by the total number of unique genes of two CHD types. We further obtained adjusted $P$ values by correcting for multiple testing using the Benjamini–Hochberg (BH) false discovery rate (FDR). Based on the matrix of Jaccard distance, which equals 1 minus the Jaccard coefficient, we performed hierarchical clustering analysis to classify the 27 CHD types into homogeneous groups using the pheatmap R package with the clustering method of ward.D. To be conservative, we reclassified the variants in CHDbase using InterVar (Version: 2.2.2) [5]. A total of 21 variants considered as “disease causing” or “likely disease causing” by the origin publications were reclassified as “benign” or “likely benign” due to high frequency in population data. We then repeated the above analyses after removing these 21 variants, and compared the results with the previous findings.

### Supplementary figures

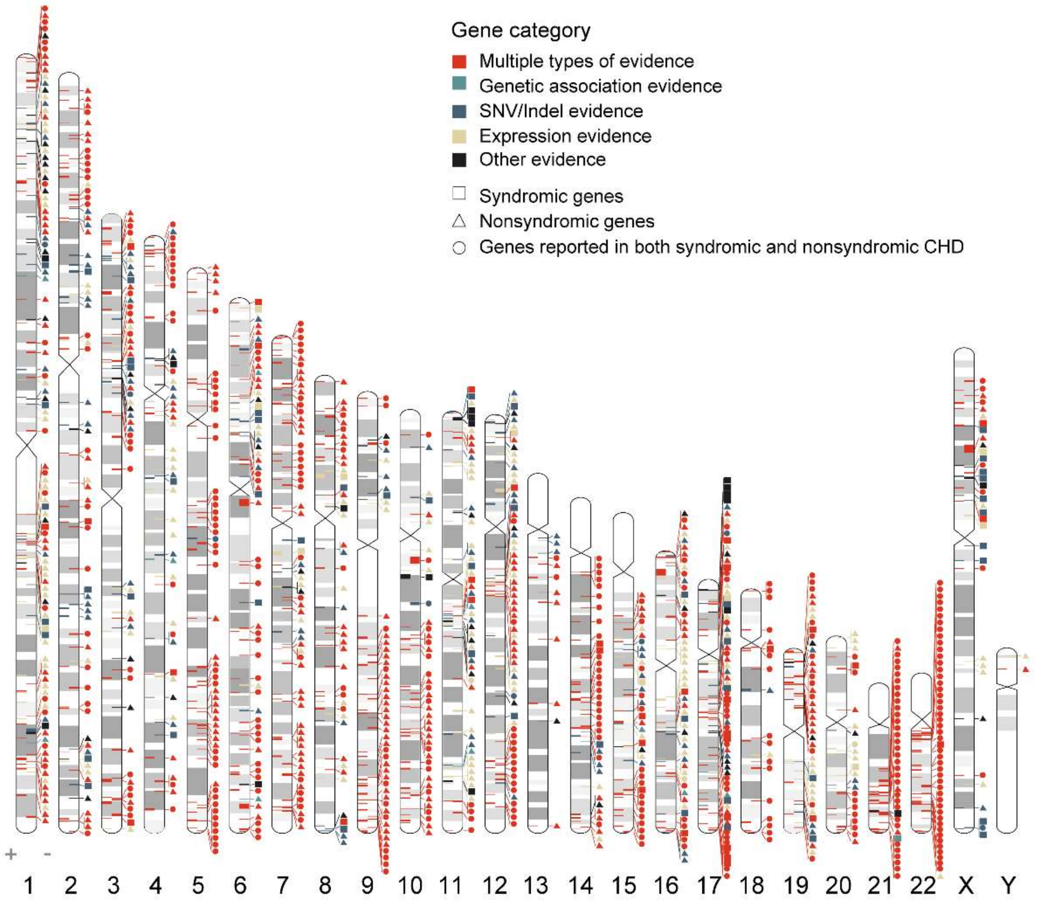


**Figure S1 Chromosome distribution for CHD-related genes**

The color of each gene represents the type of evidence supporting the association of the gene with CHD. The shape of each gene represents the category of the gene.

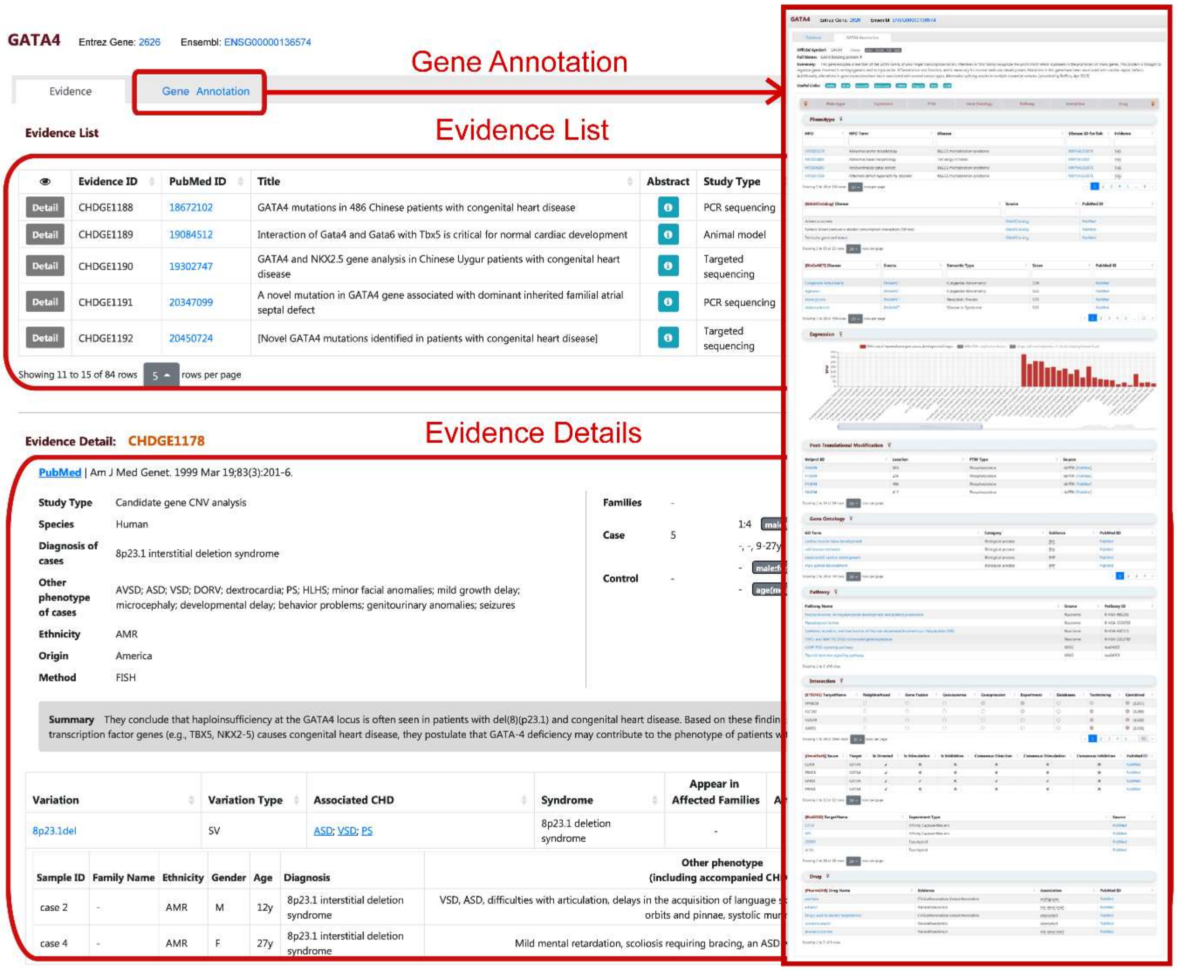


**Figure S2 Typical Gene Evidence Page and Annotation Page in CHDbase**

The Gene Evidence Page displays the evidence details at the gene level. Gene annotations can be obtained by clicking on the label "Gene Annotation" in the top left.

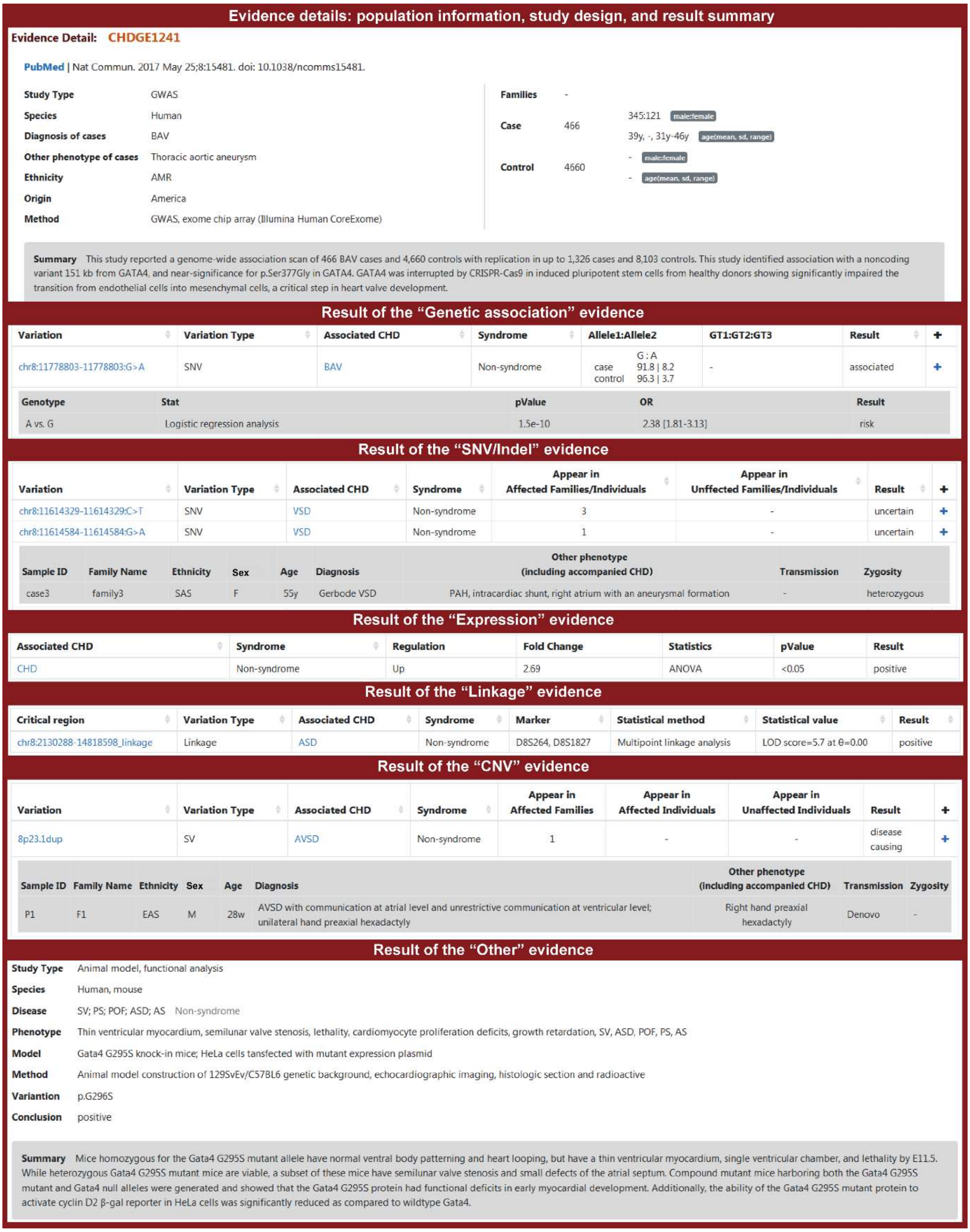


**Figure S3 Examples of evidence details at the gene level**

As shown in **Figure S2**, the evidence list is shown in the upper panel of the Gene Evidence Page, and the details of the evidence are shown in the lower panel. In the lower panel, population information, study design, and result summary are provided at the top for all types of evidence, and result details are provided in the lower part in different formats for different types of evidence.

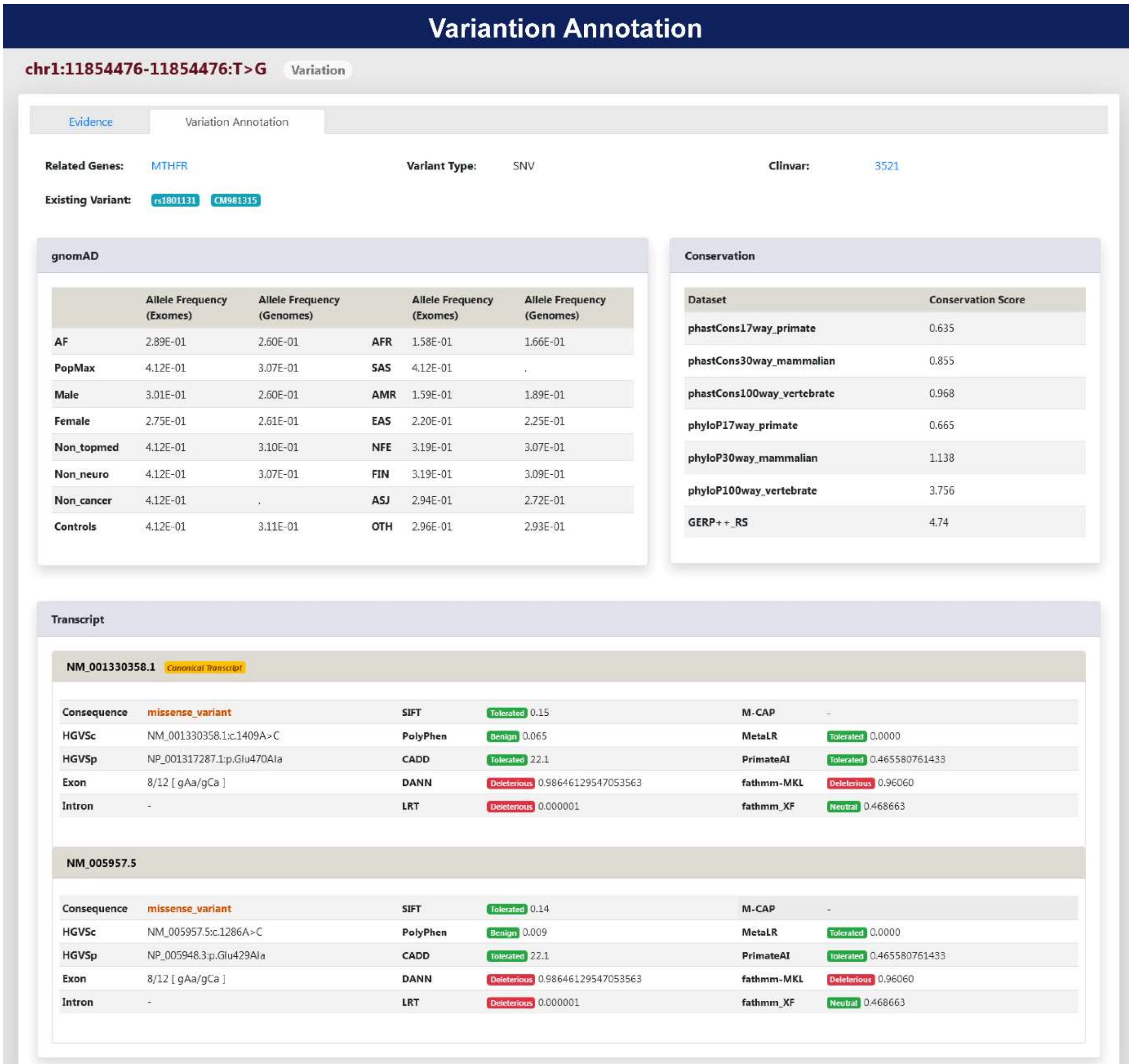


**Figure S4 Typical Variation Annotation Page in CHDbase**
